# An astrocytic basis of caloric restriction action on the brain plasticity

**DOI:** 10.1101/625871

**Authors:** Alex Plata, Alexander Popov, Pavel Denisov, Maxim Bychkov, Alexey Brazhe, Ekaterina Lyukmanova, Natalia Lazareva, Alexei Verkhratsky, Alexey Semyanov

**Affiliations:** University of Nizhny Novgorod, Gagarin ave. 23, Nizhny Novgorod, 603950, Russia; Shemyakin-Ovchinnikov Institute of Bioorganic Chemistry, Russian Academy of Sciences, Miklukho-Maklaya street 16/10, Moscow, 117997, Russia; Moscow State University, Leninskie gory 1/12, Moscow, 119234, Russia; Moscow Institute of Physics and Technology, Dolgoprudny, Moscow Region; Russia; Sechenov First Moscow State Medical University, Moscow, Russia; Faculty of Life Sciences, The University of Manchester, Manchester, M13 9PT, UK; Achucarro Center for Neuroscience, IKERBASQUE, Basque Foundation for Science, 48011 Bilbao, Spain 4

**Keywords:** astrocyte, calcium, glutamate transporters, spillover, LTP, caloric restriction

## Abstract

One month of calorically restricted diet (CR) induced morphological plasticity of astrocytes in the *stratum (str.) radiatum* of hippocampal CA1 in three-months old mice: the volume fraction of distal perisynaptic astrocytic processes increased whereas the number of gap-junction coupled astrocytes decreased. The uncoupling was not associated with a decrease in the expression of connexin 43. Uncoupling and morphological remodeling affected spontaneous Ca^2+^ activity in the astrocytic network: Ca^2+^ events became longer, whereas their spread was reduced. The change in the pattern of astrocytic Ca^2+^ activity may increase the spatial resolution of the information encoding in the astroglial network. Consistent with expanded synaptic enwrapping by the astroglial processes, the spillover of synaptically released K^+^ and glutamate was diminished after CR. However, no significant changes in the expression of astrocytic glutamate transporter (GLT-1/EAAT2) were observed, although the level of glutamine synthetase was decreased. Glutamate uptake is known to regulate the synaptic plasticity. Indeed, the magnitude of long-term potentiation (LTP) in the glutamatergic CA3-CA1 synapses was significantly enhanced after CR. Our findings highlight an astroglial basis for improved learning and memory reported in various species subjected to CR.

## Introduction

Changes in demographic profile and rapid growth of elderly population during last century arguably represent the most dramatic metamorphosis of human society since the emergence of *Homo Sapiens*; the prevalence of aged population in conjunction with destroyed family canvass and technological progress are fundamental challenges that define the future of humanity. The notion of smart aging (which assumes substantial prolongation of normal human activities at the advanced age) starts to emerge worldwide (Song et al., 2018). Food consumption and dieting, as well as lifestyle and physical exercises, are essential determinants of lifespan as well as cognitive capabilities (Mattson, 2012). Prominent positive effect of caloric restriction on the lifespan of rats was discovered in 1935 (McCay et al., 1935), and have been confirmed since for several species; the restrictive dieting may also have some beneficial effects on senescent-dependent pathology in primates and humans (Fontana et al., 2010; Mattison et al., 2017; Mattson, 2012; Ngandu et al., 2015; Redman et al., 2008). Although several underlying mechanisms have been considered (e.g., reduced oxidative damage, tamed inflammation or accumulation of ketones (Maalouf et al., 2009; Masoro, 2009; Veech et al., 2017)) none become universally acknowledged. Restrictive dieting affects the brain aging; low calories intake has been noted to exert neuroprotection, retard age-dependent cognitive decline and decrease the incidence of neurodegenerative diseases (Maalouf et al., 2009). Again, several underlying mechanisms were proposed, and yet little systematic studies of cellular physiology of neural cells from CR exposed animals have been executed. Neuroprotection in the CNS is assigned to glial cells; with astrocytes being particularly important for supporting neuronal networks.

The aging of the brain with associated cognitive decline and neurodegenerative diseases (including Alzheimer’s disease), is, to a large extent, defined by the lifestyle. Insufficient cognitive engagement, lack of exercise and excessive food intake promote, whereas cognitive stimulation, physical activity and dieting delay age-dependent cognitive deterioration (Mattson, 2015; Ngandu et al., 2015). Energy intake and energy balance are critically important for the brain, which requires high glucose consumption to maintain ionic gradients and hence excitability and synaptic transmission (Magistretti, 2009). How restricting caloric intake affects the physiology of the neural cells remains largely unexplored. Low-calorie dieting, similarly to other modified food intake programmes, impacts neuronal circuitry of the hypothalamus (Garcia-Caceres et al., 2019; Zeltser et al., 2012), which regulates energy balance. There is also evidence indicating that the CR affects the physiology of neurons in other brain regions. Exposure to CR increases memory and boosts brain plasticity (Kuhla et al., 2013; Murphy et al., 2014). At a cellular level CR prevents an age-dependent decline in hippocampal synaptic plasticity (Okada et al., 2003), enhances neurogenesis (Fontan-Lozano et al., 2008) and improves neuroplasticity in visual cortex (Spolidoro et al., 2011), through increasing intracortical inhibition by up-regulation of GABA synthesis and neuronal GABA content (Yang et al., 2016). There are also claims that CR increases neuronal metabolism and preserves neuronal activity in the aged brain (Lin et al., 2014). At the same time restricting calorie intake by 30% for four years in lemurid primate *Microcebus murinus* led to a significant life prolongation (from 6.4 to 9.6 years) paralleled with accelerated loss of grey matter (Pifferi et al., 2018). Whether CR causes such brain shrinkage, or it naturally occurs with aging beyond the normal lifespan remains to be established.

The functional activity of neuronal networks, as well as adaptive and life-long neuroprotection, is supported by neuroglia; in particular astrocytes, the homeostatic cells of the central nervous system, safeguard brain homeostasis at all levels of organization from molecular to organ (Verkhratsky and Nedergaard, 2018). Astrocyte distal processes enwrap synapses creating an astroglial cradle that regulates synaptogenesis, synaptic maturation, synaptic isolation, maintenance, and extinction (Verkhratsky and Nedergaard, 2014). Astrocytes are also intimately involved in supporting neuronal metabolism and in regulating neurotransmitter balance (Pellerin and Magistretti, 2012; Verkhratsky and Nedergaard, 2018). How the restriction of calorie intake affects astrocytes is virtually unknown; a sporadic report has demonstrated that chronic CR causes a decrease in the size of astrocytes in mice of 19 to 24 months of age when compared to the *ad libitum* fed controls (Castiglioni Jr et al., 1991). Here we tested the effect of CR on astrocytes and brain plasticity in the younger age group.

## Results

Two-month-old mice were subjected to CR for one month and then compared to the same age animals receiving food *ad libitum* (control). CR mice showed a small weight loss, while the control group showed a small weight increase. After a month, the control mice gained weight to 114 ± 3 % (*n* = 22), while the animals in CR group lost weight to 85 ± 4 % (*n* = 14; *p* < 0.001, two sample *t*-test; Fig. S1).

### CR increases astrocytic perisynaptic coverage

To reveal the morphology of individual protoplasmic astrocytes, the cells were filled with a fluorescent tracer Alexa Fluor 594 through a patch-pipette and imaged with two-photon laser scanning microscopy in CA1 *stratum (str.) radiatum* of hippocampal slices. Sholl analysis did not show any significant difference in the number of optically resolved astrocytic branches at different distances from the soma between CR and control mice (control: *n* = 7; SE: *n* = 7; *F*(_1,6_) = 0.267, *p* = 0.624 for the effect of CR, partial *η*^2^ = 0.043, Sphericity assumed, two-way repeated measures ANOVA; Fig. 1a,b). The Sholl analysis cannot reflect changes in fine perisynaptic astrocytic leaflets which are beyond the resolution of diffraction-limited optical microscopy (Khakh and Sofroniew, 2015; Semyanov, 2019). However, the volume fraction (VF) of perisynaptic leaflets can be indirectly measured as fluorescence ratio of unresolved processes area to the astrocyte soma (Medvedev et al., 2014; Plata et al., 2018). This type of analysis assumes that soma fluorescence reflects 100% of astrocyte space occupancy, while the fluorescence of unresolved area is proportional to the VF of astrocyte processes in this area (Fig. 1c). This analysis revealed that VF of perisynaptic leaflets was significantly increased in astrocytes of CR mice (control: 3.2 ± 0.4 %, *n* = 7; CR: 4.5 ± 0.4 %, *n* = 7; *p* < 0.001, two sample *t*-test; Fig. 1d).

**Fig. 1.**
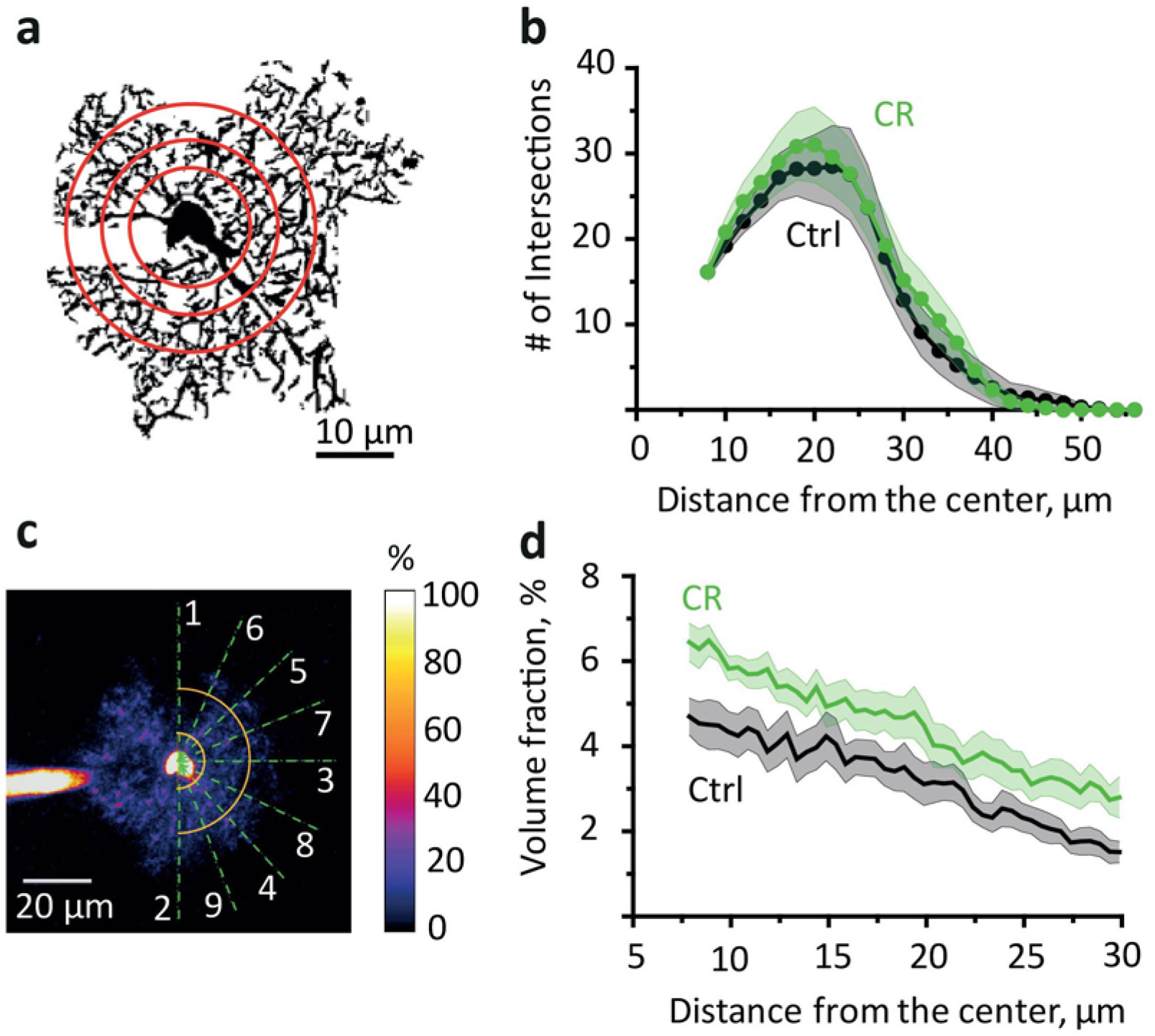
CR does not affect astrocytic branches but increases the VF of perisynaptic astrocytic leaflets. **a.** A mask of astrocytic branches used for Sholl analysis (red circles) was obtained from a maximal projection of z-stack of fluorescence images of the astrocyte loaded with 50 μM Alexa Fluor 594 through patch pipette. **b.** The summary data for the number of intersections of circles drawn around the center of astrocyte soma with astrocytic branches. Green – CR, black – control (Ctrl). **c.** Reconstruction of fluorescence profiles across an astrocyte to estimate volume fraction (VF) of unresolved processes. Dashed lines indicate the places where the profiles were obtained. The profiles included the soma and the area or unresolved processes. The large local increases in fluorescence corresponding to astrocytic branches were cut out. Yellow semicircles indicate the area analyzed for this astrocyte to avoid an effect of astrocyte domain asymmetry. **d.** The summary data showing estimated VF of unresolved astrocytic processes outside of soma. The data presented as mean ± SEM.

### CR disrupts astroglial gap-junction coupling

Next, we analyzed the effect of CR on the astrocytic syncytial network. The density of astrocytes labeled with astrocyte-specific marker sulforhodamine 101 was similar in the control and CR mice (control: 1.8 ± 0.4 astrocytes per 100^2^ µm^2^, *n* = 6; CR: 1.8 ± 0.4 astrocytes per 100^2^ µm^2^, *n* = 13; *p* = 0.93, two sample *t*-test; Fig. S2). The cell coupling within the astrocytic network was analyzed with the method based on the diffusion of the intercellular dye through the gap-junctions (Henneberger et al., 2010; Plata et al., 2018). The number of coupled astrocytes was significantly lower in CR than in the control mice (control: 12.6 ± 1.9, *n* = 5; CR: 4.2 ± 1.3, *n* = 5; *p* = 0.003, two sample *t*-test; Fig. 2a,b). The dye diffusion technique also permits to estimate the strength of astrocytic coupling. The fluorescence measured from the soma of coupled astrocytes decays exponentially with the distance from the patched astrocyte (Fig. 2c, note semi-logarithmic scale). The length constant did not show a significant difference between two groups of animals (control: 19.9 ± 1.9 µm, *n* = 5; CR: 26.1 ± 4.3 µm, *n* = 5; *p* = 0.12, two sample *t*-test; Fig. 2d). This finding indicates that despite a decrease in the number of coupled astrocytes, the strength of remaining gap-junction connections is unaffected. The astrocyte uncoupling may be linked to morphological remodeling of coupled processes or to changes in expression of connexins responsible for gap-junction formation. To address this possibility, we performed Western blotting and did not detect a significant difference in the expression of astrocyte-specific connexin 43 (protein level normalised to β-actin in control: 1.09 ± 0.19, *n* = 5; in CR: 1.03 ± 0.35, *n* = 5; *p* = 0.89, two sample *t*-test; Fig. 2e,f). Thus, astrocytic coupling changes independently of connexin 43 expression in CR mice.

**Fig. 2.**
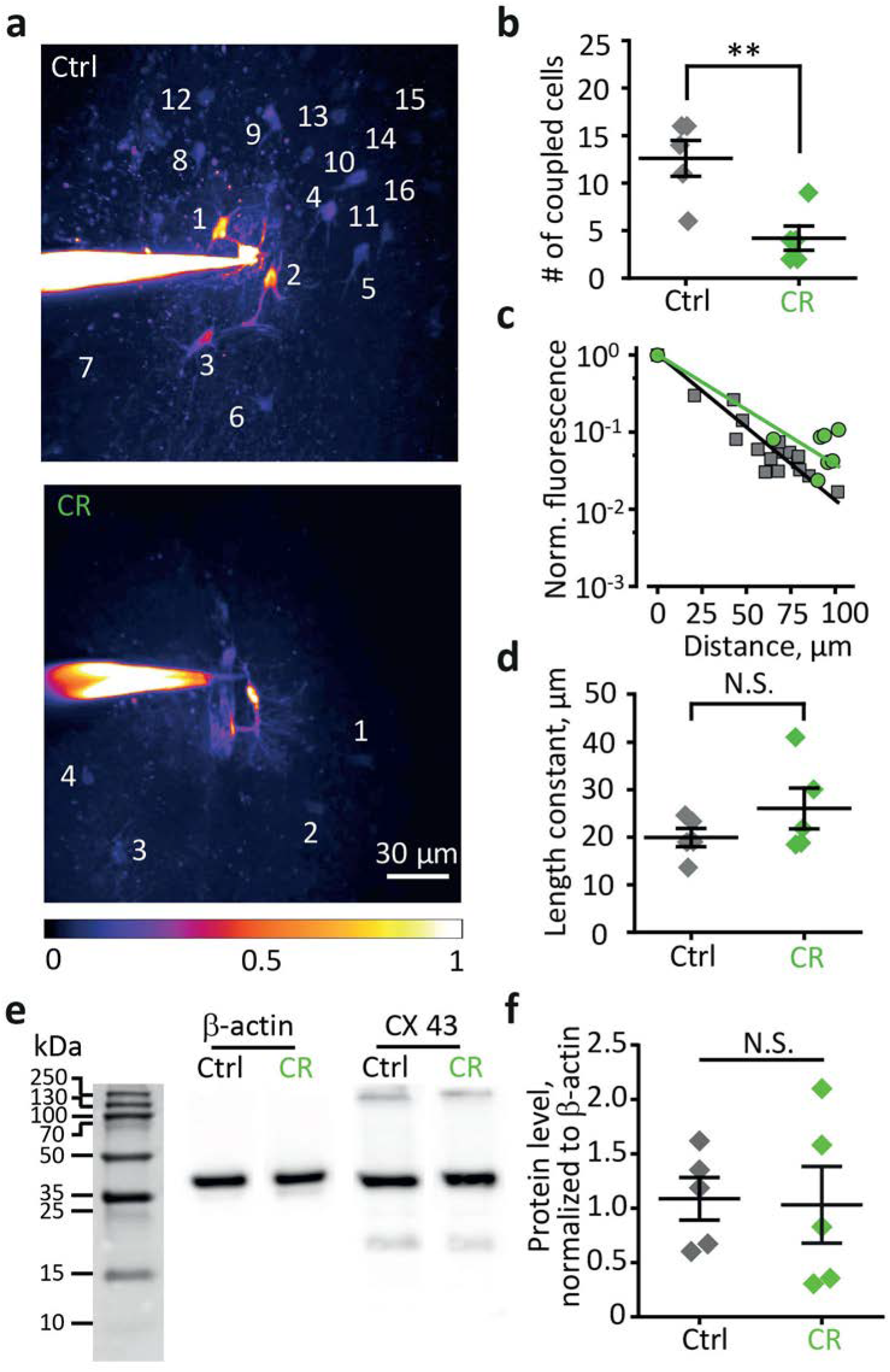
CR reduces cell coupling in the astrocytic network but does not affect the coupling strength. **a.** Fluorescence image of an astrocyte stained with 50 µM Alexa Fluor 594 through patch pipette in control (*top*) and CR mice (*bottom*). The dye diffuses through gap-junctions thus staining coupled astrocytes (numbered). The number of stained astrocytes decreased in CR. The image also shows the distance-dependent decrease in the somatic fluorescence of coupled astrocytes. The scale shows color-coding of fluorescence normalized to the fluorescence of patched astrocyte soma. **b.** The summary data showing the number of coupled astrocytes in control (grey diamonds) and CR mice (green diamonds). **c.** The decay of fluorescence in somata of coupled astrocytes with distance from the astrocyte loaded with Alexa Fluor 594. The slope of the linear fit in semi-logarithmic scale determines the length constant. Grey squares – control and grin circles – CR. **d.** The summary data showing the length constant in control (grey diamonds) and in CR mice (green diamonds). **e.** Representative western blots of the mouse hippocampus homogenates stained by antibodies against β-actin and connexin 43 (CX 43). **f.** Normalized to β-actin optical densities of the western blot bands for CX43. Grey diamonds – control, green diamonds – CR mice. The data are presented as mean ± SEM; N.S. *p* > 0.05; * *p* < 0.05; two-sample *t*-test.

### CR modifies spatiotemporal properties of spontaneous Ca^2+^ events in the astrocytic network

Morphological remodeling and gap-junction uncoupling can affect Ca^2+^ activity in the astroglial network (Plata et al., 2018; Semyanov, 2019; Wu et al., 2019). We performed confocal imaging of astrocytes stained with membrane-permeable Ca^2+^ dye, Oregon Green 488 BAPTA-1 AM (OGB, Fig. 3a). The areas of Ca^2+^ events were identified in each frame, and individual events were reconstructed in three-dimensions (x-y-time, Fig. 3b) (Wu et al., 2014). Only Ca^2+^ events longer than 2 s were analyzed to avoid the contribution of fast spontaneous Ca^2+^ transients in neurons that can also be stained with OGB (Monai et al., 2016; Plata et al., 2018). The frequency of detected Ca^2+^ events was not significantly different in CR mice (control: 0.33 ± 0.08 events/astrocyte/min, *n* = 11; CR: 0.37 ± 0.10 events/astrocyte/min, *n* = 10; p = 0.77, two sample *t*-test; Fig. 3c). However, more Ca^2+^ events with longer duration appeared in the distribution (*p* = 0.042, Mann-Whitney test with control; Fig. 3b,d). The proportion of Ca^2+^ events with the larger size, in contrary, decreased in CR mice when compared to the control group (*p* = 0.006, Mann-Whitney test with control, Fig. 3b,e). This redistribution of the Ca^2+^ event properties in CR can be best seen at the relationship between the duration and size of the events (Fig. 3f). Such changes in Ca^2+^ activity arguably reflect cell uncoupling in the astrocytic network and the inability of Ca^2+^ events to propagate through the gap-junctions to the neighboring cells (Fujii et al., 2017). The integral of astrocytic Ca^2+^ activity was unchanged in CR (*p* = 0.27, Mann-Whitney test with control, Fig. 3g).

**Fig. 3.**
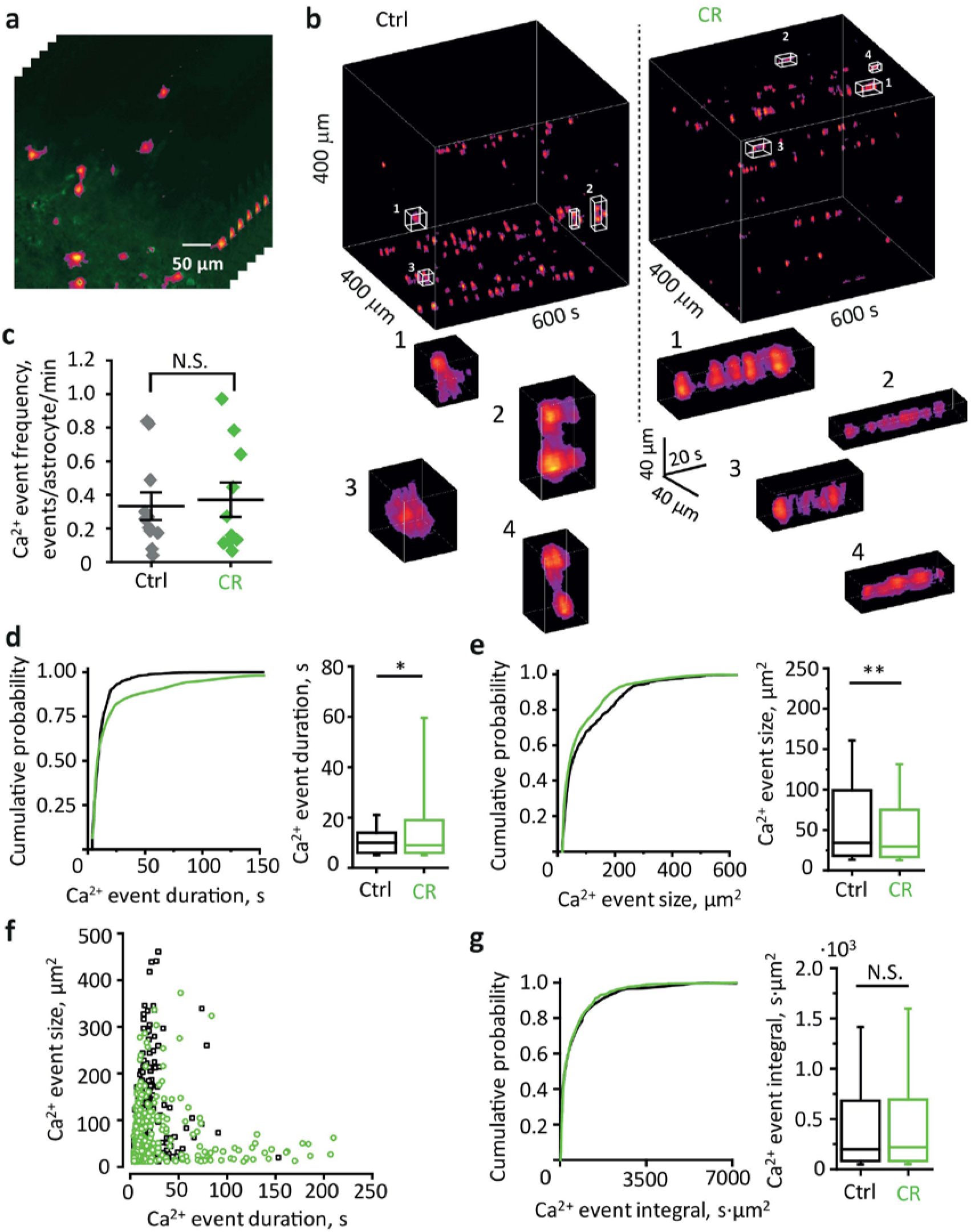
CR decreases size but increases the duration of astrocytic Ca^2+^ events. **a.** z-stack of fluorescence images in green with superimposed color-coded relative fluorescence changes. **b.** Individual Ca^2+^ events reconstructed in three-dimensions: x-y-time. *Left*, control; *right*, CR mice. *Top*, population Ca^2+^ activity in the astrocytic network in the 400 × 400 µm^2^ square during 600 s. *Bottom*, representative examples of Ca^2+^ events marked in the population activity above. Note enlargement of Ca^2+^ events due to their spread in the x-y plane in control mice; and due to their prolongation in the time axis in CR mice. **c.** The summary plot of Ca^2+^ event frequency normalized to the number of astrocytes identified within the recording area. No significant difference observed between control (grey diamonds) and CR (green diamonds) mice. **d.** Cumulative probability (*left*) and bar-and-whisker plot (*right*) for durations of all Ca^2+^ events in control (*n* = 669 events, *N* = 11 mice) and in CR (*n* = 440 events, *N* = 10 mice). The result demonstrates the appearance of long-lasting Ca^2+^ events in CR mice. **e.** Cumulative probability (*left*) and bar-and-whisker plot (*right*) for sizes of all Ca^2+^ events in control and in CR mice. The result demonstrates the loss of large size Ca^2+^ events in CR mice. **f.** A graph showing all Ca^2+^ events in duration-size space. The enlargement of Ca^2+^ events occurs due to their spread in an x-y plane without prolongation in control mice (black squares) and due to the prolongation without an increase in the spread in CR mice (green circles). **g.** Cumulative probability (*left*) and bar-and-whisker plot (*right*) for integrals (x-y-time) of all Ca^2+^ events in control and in CR mice. The result demonstrates that Ca^2+^ activity remodeling does not affect the integral of Ca^2+^ events in CR mice. The data at panel c. are presented as mean ± SEM; N.S. *p* > 0.05, two sample *t*-test. The data at panels d., e., g. presented as median, quartiles and 10-90 whisker range; N.S. *p* > 0.05, * *p* < 0.05, * *p* < 0.01, Mann-Whitney test.

### CR enhances activity-dependent K^+^ release but reduces K^+^ spillover

Increased VF of perisynaptic astrocytic leaflets may modify major homeostatic functions including K^+^ clearance. During synaptic transmission, K^+^ is mostly released through postsynaptic ionotropic glutamate receptors, predominantly of NMDA type (Ge and Duan, 2007; Shih et al., 2013; Sibille et al., 2014). Thus, astrocytic K^+^ current (I_K_) in astrocyte can provide a readout of both postsynaptic response (the I_K_ amplitude) and astrocytic K^+^ clearance (the I_K_ decay time [τ_decay_ I_K_], Fig. 4a). We recorded I_K_ in response to stimulation of Schaffer collaterals in the absence of synaptic receptor blockers in CA1 *str.radiatum* astrocytes. The early part of I_K_ overlapped with fast glutamate transporter current (I_GluT_); therefore, the I_K_ amplitude was measured as a maximal current 200 ms after the last stimulus. The τ_decay_ I_K_ was determined by the exponential fit of the I_K_ current. To estimate activity-dependent changes in I_K_, the response to the fifth stimulus (I_K_(5)) in the train of five stimuli (5 × 50 Hz) was isolated and compared to the response to a single stimulus (I_K_(1), Fig. 4b). We observed a decrease in five-pulse ratio of I_K_ in the astrocytes of control mice, whereas in CR mice it was increased (I_K_(5)/I_K_(1) in control: 0.81 ± 0.07, *n* = 10; *p* = 0.03, one sample *t*-test; in CR: 1.21 ± 0.03, *n* = 11; *p* <0.001, one sample *t*-test; *p* < 0.001, two sample *t*-test between control and CR; Fig. 4c,d). Notably, the five-pulse ratio of I_K_ was not significantly different between control and CR mice in the presence of NMDA, AMPA and GABA_A_ receptors blockers (I_K_(5)/I_K_(1) in control: 0.89 ± 0.05, *n* = 7; in CR: 1.02 ± 0.09, *n* = 7; *p* = 0.23, two sample *t*-test; Fig. S3a,b). Because GABA_A_ receptors are not K^+^ permeable, this result suggests that CR promotes activity-dependent facilitation at glutamate synapses.

**Fig. 4.**
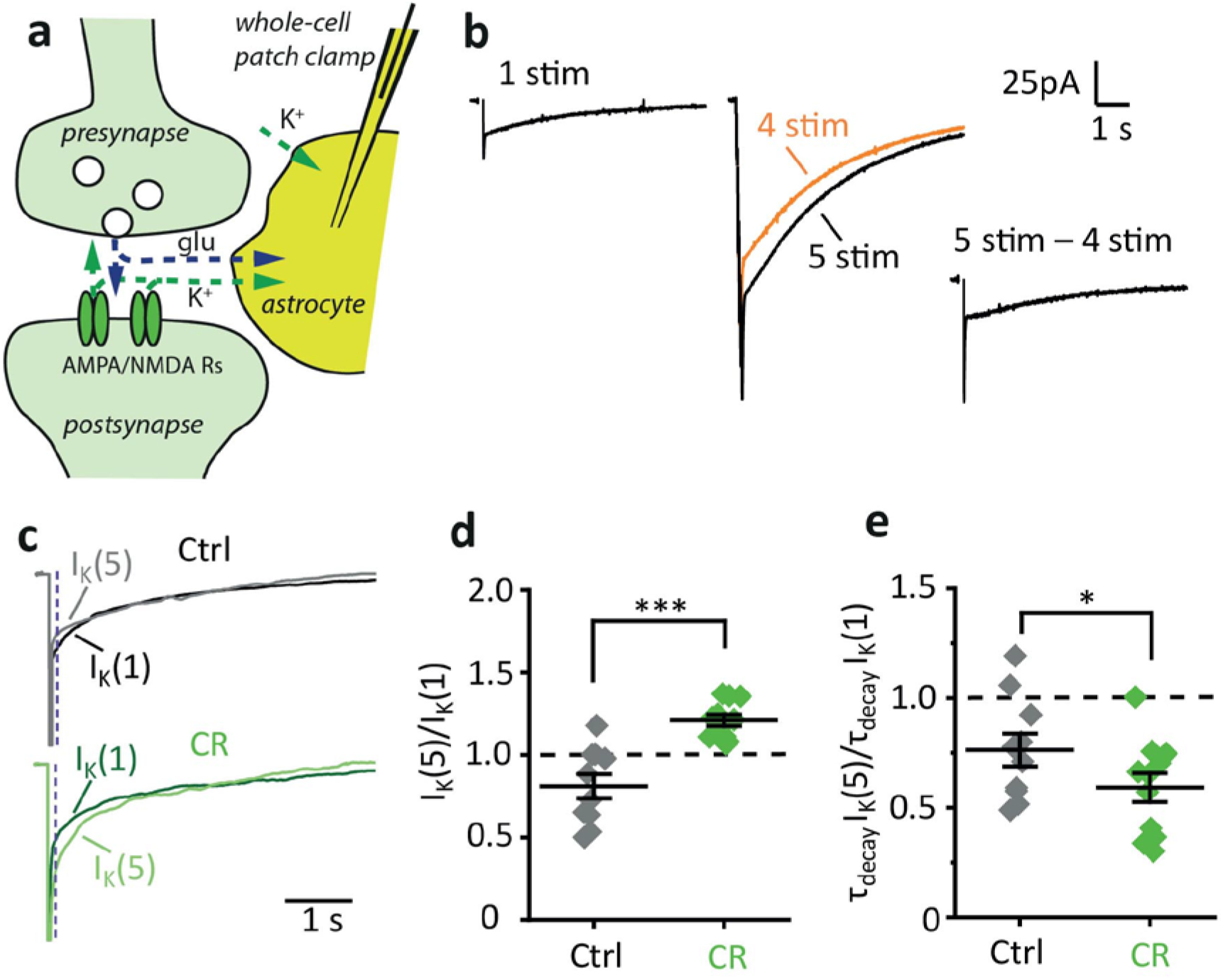
CR enhances activity-dependent K^+^ release but reduces K^+^ spillover. **a.** A schematic of the main mechanisms of the synaptically induced current in astrocyte. Glutamate released by perisynaptic terminal triggers transient current mediated by astrocytic transporters (I_GluT_). This glutamate also activates postsynaptic ionotropic receptors (AMPA and NMDA type) which are permeable for K^+^. K^+^ efflux through postsynaptic glutamate receptors is responsible for most of I_K_ in astrocyte (small amount of I_K_ is mediated by other mechanisms such as action potentials). **b.** An experimental protocol to estimate activity-dependent changes in I_K_. *Left*, the astrocytic current induced by a single stimulus. Fast I_GluT_ is followed by slow I_K_. *Middle*, the astrocytic current induced by 5 stimuli (black trace) superimposed by the astrocytic current induced by 4 stimuli (orange trace). *Right*, the current to fifth stimulus isolated by subtraction of the current to 4 stimuli (at 50 Hz) from the current to 5 stimuli (at 50 Hz). **c.** Representative currents to a single stimulus (dark traces) and the fifth stimulus (light traces) in control (grey) and CR mice (green). The dashed line shows where the measurement of I_K_(1) and I_K_(5) amplitudes and decay times were taken to avoid the interference of I_GluT_. **d.** The summary plot showing an increase in the five pulse ratio – i.e., I_K_(5)/I_K_(1), in CR mice. Grey diamonds – control, green diamonds – CR mice. **e.** The summary plot showing a decrease in the five pulse ratio of decay time – i.e., τ_decay_I_K_(5)/τ_decay_ I_K_(1), in CR mice. Grey diamonds – control, green diamonds – CR mice. The data are presented as mean ± SEM; * *p* < 0.05; *** *p* < 0.001; two sample *t*-test.

The five-pulse ratio of τ_decay_ I_K_ decreased both in CR and control mice, but the decrease was more profound in CR animals (τ_decay_ I_K_(5)/τ_decay_ I_K_(1) in control: 0.76 ± 0.08, *n* = 10, *p* = 0.01, one-sample *t*-test; in CR: 0.59 ± 0.06, *n* = 11, *p* < 0.001 one-sample *t*-test; *p* = 0.042, two-sample *t*-test between control and CR; Fig. 4e). This finding indicates enhanced K^+^ clearance and is consistent with increased astrocytic coverage of synapses in CR mice. Again, the five-pulse ratio of τ_decay_ I_K_ did not significantly differ between CR and control mice in the presence of synaptic receptor blockers (τ_decay_ I_K_(5)/τ_decay_ I_K_(1) in control: 1.01 ± 0.30, *n* = 8; in CR: 0.89 ± 0.27, *n* = 8; *p* = 0.39, two sample *t*-test; Fig. S3a,c).

### CR reduces glutamate spillover

Next, we recorded I_GluT_ in astrocytes. Because K^+^ accumulation in the synaptic cleft affects both presynaptic release probability and glutamate uptake efficiency (Ge and Duan, 2007; Lebedeva et al., 2018; Shih et al., 2013), further recordings were performed in the presence of NMDA, AMPA and GABA_A_ receptor blockers (Fig. 5a). The residual I_K_ was pharmacologically isolated with 100 µM DL-TBOA, a glutamate transporter blocker, and subtracted from the combined synaptically induced current to reveal pure I_GluT_ (Lebedeva et al., 2018; Scimemi and Diamond, 2013). The response to the fifth stimulus (I_GluT_(5)) in the train (5 × 50 Hz) was compared to the response to a single stimulus (I_GluT_(1), Fig. 5b). The five-pulse ratio of I_GluT_ was consistent with activity dependent-facilitation of presynaptic glutamate release and was not significantly different between astrocytes of CR and control groups (I_GluT_(5)/I_GluT_(1) in control: 1.56 ± 0.25, n = 8, p = 0.03, one sample *t*-test; in CR: 1.43 ± 0.15, *n* = 8, *p* = 0.012 one-sample *t*-test; p = 0.69, two sample *t*-test between control and CR; Fig. 5c,d). The five-pulse ratio of τ_decay_ I_GluT_ in control mice was larger than 1:1, reflecting glutamate spillover (τ_decay_ I_GluT_(5)/τ_decay_ I_GluT_(1): 1.28 ± 0.06, *n* = 8, *p* = 0.002, one sample *t*-test). In contrast, the five-pulse ratio of τ_decay_ I_GluT_ in CR mice was not significantly different from 1:1 (τ_decay_ I_GluT_(5)/τ_decay_ I_GluT_(1): 0.97 ± 0.12, *n* = 8, *p* = 0.8, one sample *t*-test; *p* = 0.02, two-sample *t*-test for difference with control group; Fig. 5e). Thus, increased VF of perisynaptic leaflets correlates with reduced glutamate spillover in the hippocampus of CR animals. An alternative explanation for reduced glutamate spillover would be upregulation of glutamate transporters; a previous report suggested an increase of hippocampal glutamate uptake and glutamine synthetase activity in rats subjected to 12 weeks of CR (Ribeiro et al., 2009). However, we did not find a significant difference in the astroglia specific glutamate transporter 1 (GLT1/EAAT2) expression quantified with Western blotting (protein level normalised to β-actin in control: 1.02 ± 0.14, *n* = 5; in CR: 1.27 ± 0.25, *n* = 5; *p* = 0.43, two sample *t*-test; Fig. 6a,b). At the same time the level of glutamine synthetase was significantly reduced in CR mice (protein level normalised to β-actin in control: 0.98 ± 0.11, *n* = 5; in CR: 0.60 ± 0.08, *n* = 5; *p* = 0.03, two sample *t*-test; Fig. 6b,c).

**Fig. 5.**
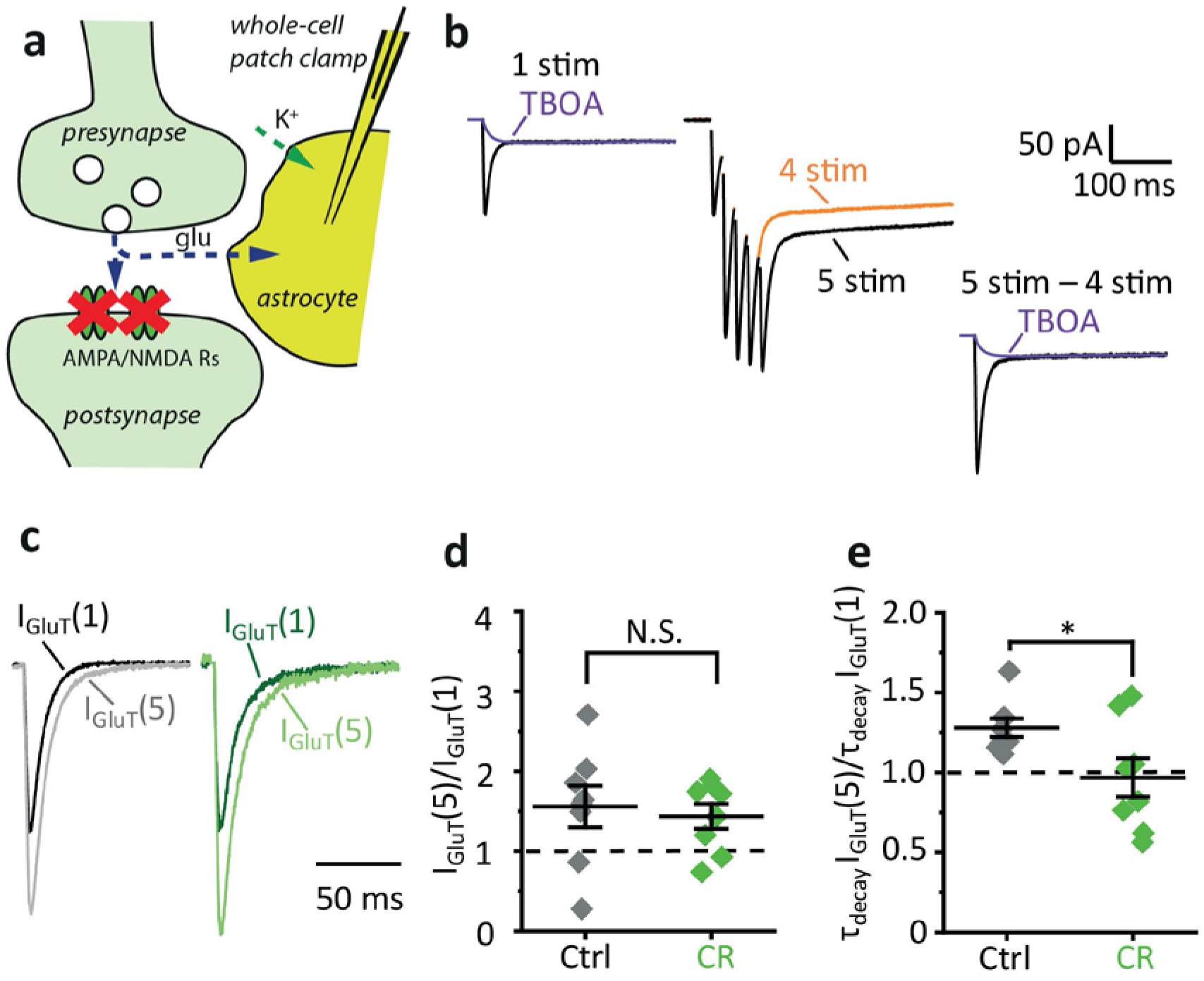
CR reduces glutamate spillover. **a.** A scheme illustrating the effect of ionotropic receptor blockers on synaptically induced astrocytic current. Glutamate released by perisynaptic terminal triggers transient current mediated by astrocytic transporters (I_GluT_) but does not trigger K^+^ efflux through postsynaptic AMPA and NMDA receptors. Only a small I_K_ is mediated by other mechanisms such as action potentials is recorded in astrocyte. **b.** An experimental protocol to estimate activity-dependent changes in I_GluT_. *Left*, astrocytic currents induced by a single stimulus in the presence of ionotropic receptor blockers with (violet trace) and without (black trace) glutamate transporter blocker (TBOA). *Middle*, the astrocytic current induced by 5 stimuli (black trace) superimposed by the astrocytic current induced by 4 stimuli (orange trace). *Right*, the current to fifth stimulus isolated by subtraction of the current to 4 stimuli (at 50 Hz) from the current to 5 stimuli (at 50 Hz) and tail-fit by the current to single stimulus in the presence of TBOA. I_GluT_(1) and I_GluT_(5) were then isolated by subtraction of TBOA-insensitive current. **c.** Representative isolated I_GluT_(1)(dark traces) and I_GluT_(5)(light traces) in control (grey) and CR mice (green). **d.** The summary plot showing similar five pulse ratio – i.e., I_GluT_(5)/I_GluT_(1), in control and CR mice. Grey diamonds – control, green diamonds – CR mice. **e.** The summary plot showing a decrease in the five pulse ratio of decay time – i.e., τ_decay_ I_GluT_(5)/τ_decay_ I_GluT_(1), in CR mice. Grey diamonds – control, green diamonds – CR mice. The data are presented as mean ± SEM; N.S. *p* > 0.05; * *p* < 0.05; two-sample *t*-test.

**Fig. 6.**
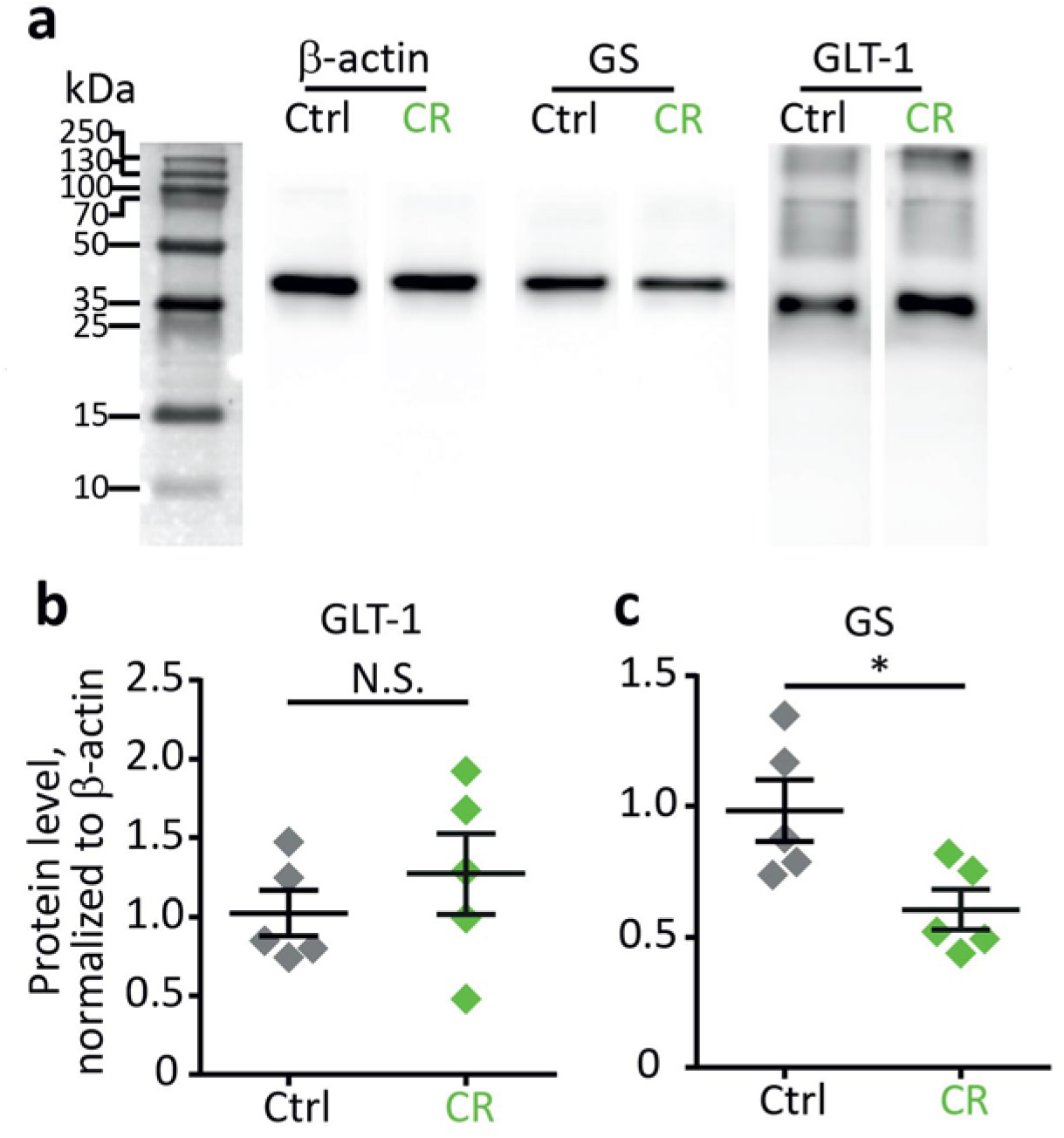
Western blot analysis of CR influence on the expression of glutamine synthetase (GS) and glutamate transporter 1 (GLT-1). **a.** Representative blots of the mouse hippocampus homogenates stained by antibodies against β-actin, GS and GLT-1. **b.** Normalized to β-actin optical densities of the western blot bands for GLT-1. Grey diamonds – control, green diamonds – CR mice. **c.** Same as b. for glutamine synthetase (GS). The data are presented as mean ± SEM; N.S. *p* > 0.05; * *p* < 0.05; two-sample *t*-test.

### CR enhances LTP in the hippocampus

Glutamate uptake regulates the extent of perisynaptic and extrasynaptic receptor activation during synaptic transmission and affects synaptic plasticity (Kullmann and Asztely, 1998; Rusakov and Kullmann, 1998). The density of glutamate transporters in the vicinity of the synapse determines the spatiotemporal properties of glutamate concentration and therefore glutamate access to pre-, post- and extrasynaptic receptors (Romanos et al., 2019). Glutamate transport consequently has been implicated in the polarity and magnitude of synaptic plasticity (Valtcheva and Venance, 2019). Downregulation of glutamate transporters leads to a reduction of the magnitude of long-term potentiation (LTP) (Katagiri et al., 2001; Scimemi et al., 2009; Wang et al., 2006) and promotes instead the long-term depression (LTD) (Brasnjo and Otis, 2001; Massey et al., 2004; Wong et al., 2007). CR increases the degree of perisynaptic coverage by astrocytic processes, hence increasing available glutamate transporters with subsequent reduction of glutamate spillover. Therefore, we hypothesized that LTP might be enhanced after CR.

We performed field potential recordings in CA1 *str.radiatum* in response to extracellular stimulation of Schaffer collaterals. The relationships of presynaptic volley (PrV) versus stimulus and field excitatory postsynaptic potential (fEPSP) versus stimulus (input-output characteristics) were not affected by CR (Fig. 7a,b). However, we observed an enhancement of LTP induced by high-frequency stimulation (HFS) in CA3-CA1 synapses of CR mice (Fig. 7c). LTP magnitude was higher both at the early stage (first 10 min after HFS: 168 ± 9 % of baseline in control, n = 6; 204 ± 10 % of baseline in CR, *n* = 6; *p* = 0.01, two sample *t*-test; Fig. 7d) and at the later stage (40 - 50 min after HFS: 144 ± 7 % of baseline in control, *n* = 6; 162 ± 6 % of baseline in CR, *n* = 6; *p* = 0.04, two sample *t*-test; Fig. 7e). Thus plasticity of astrocytes parallels enhanced LTP in CR mice, and both may provide cellular mechanisms for enhanced learning and memory associated with this diet.

**Fig. 7.**
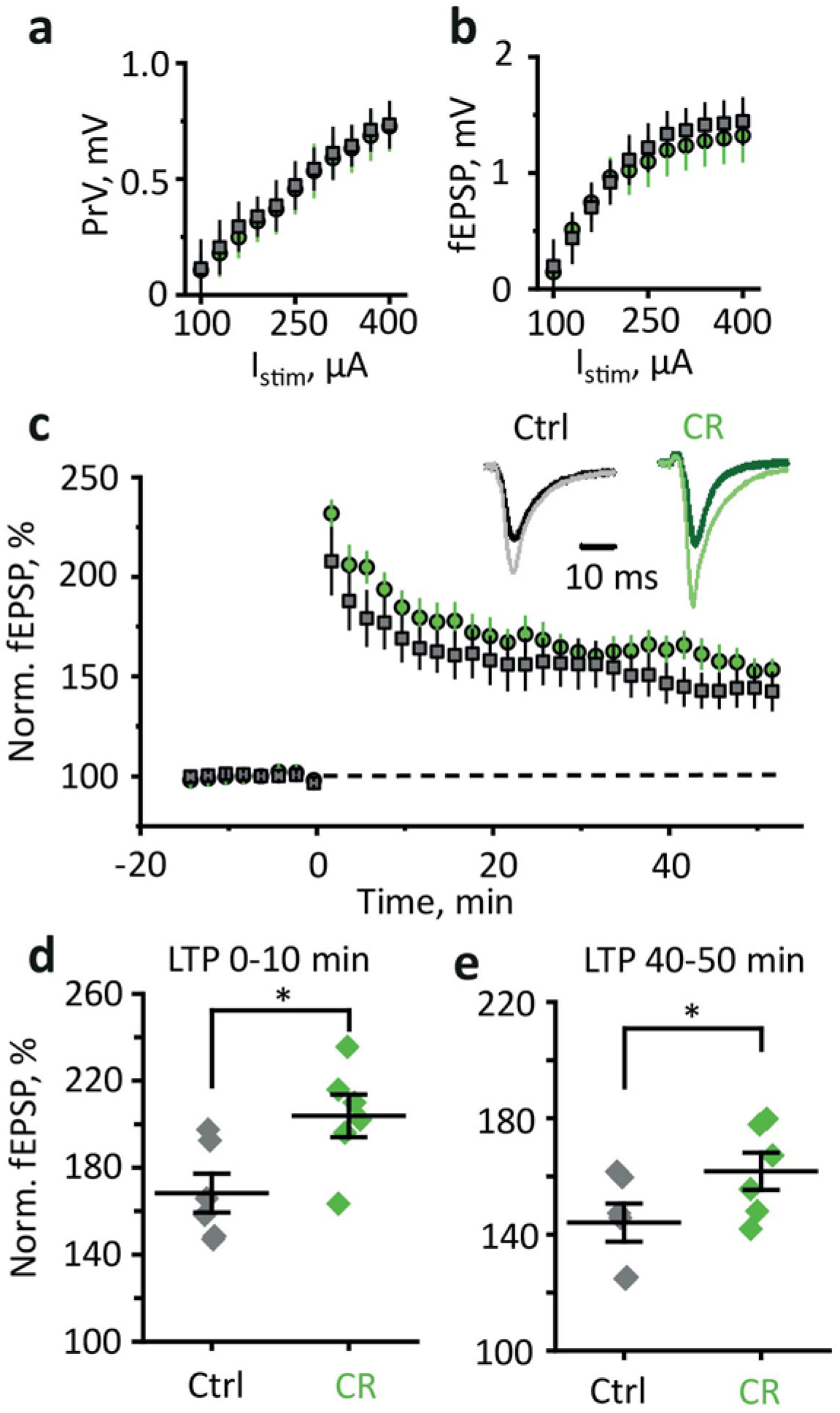
CR enhances LTP in CA3-CA1 synapses. **a,b.** Input-output relationships of PrV amplitude - stimulation current (I_stim_) (**a**) and fEPSP amplitude - I_stim_ (**b**) show no difference between control and CR mice. **c.** LTP induced by HFS in the CA1 region of the hippocampus of control (grey squares) and CR mice (green circles). *Inset*, sample fEPSPs before HFS (dark color) and 40 min after HFS (light color). Grey – control (Ctrl), green – CR. **d,e.** The summary graphs showing a significant increase in LTP in CR mice as measured by the average normalized fEPSP amplitude 0 – 10 min (**d**) and 40 – 50 min (**e**) after HFS. The data are presented as mean ± SEM; * *p* < 0.05; two-sample *t*-test.

## Discussion

The present study reports, for the first time, in-depth analysis of astroglial plasticity induced by the CR diet. We found that CR results in remodeling of protoplasmic astrocytes in the hippocampus; this astrocytic response was associated with enhanced LTP indicative of increased synaptic plasticity, which often accompanies learning associated with environmental adaptation.

Astrocytic morphological plasticity may occur at the level of individual astrocytes and at the level of the astrocytic syncytium. At the single cell level remodeling may affect astrocytic branches and astrocytic leaflets. These two types of processes are functionally distinct (Khakh and Sofroniew, 2015; Semyanov, 2019). Astrocytic branches contain organelles including Ca^2+^ stores that allow them to amplify and propagate Ca^2+^ events (Patrushev et al., 2013). Thus, remodeling of astrocytic branches would strongly affect Ca^2+^ dynamics in individual astrocytes. Atrophy of astrocytic branches and reduced spontaneous Ca^2+^ activity in astrocytes has been reported in epilepsy (Plata et al., 2018). Astrocytic branches can be resolved with two-photon microscopy in the hippocampus. Even the thinnest terminal branches (also termed branchlets) are about 1 µm in diameter, which is still within the resolution of diffraction-limited systems (Gavrilov et al., 2018; Patrushev et al., 2013). Hence, we performed Sholl analysis of astrocytic branches which did not reveal any significant changes in their number after CR.

Astrocytic leaflets are flat perisynaptic processes devoid of organelles (Fernandez et al., 1984; Gavrilov et al., 2018; Patrushev et al., 2013). Perisynaptic astrocytic leaflets (PALs) form the astroglial cradle that controls all aspects of synaptic function from synaptogenesis and synaptic maturation to synaptic maintenance, synaptic isolation, and extinction (Nedergaard and Verkhratsky, 2012; Verkhratsky and Nedergaard, 2014). Ca^2+^ dependent changes in PALs occur during synapse formation and synaptic plasticity (Perez-Alvarez et al., 2014; Schiweck et al., 2018; Tanaka et al., 2013). The decrease in astrocytic coverage of the synapses is reported in the hypothalamus of lactating rats (Oliet et al., 2001). Unlike astrocytic branches, PALs are beyond the resolution of diffraction-limited microscopy. Therefore, we used an indirect method to estimate their VF as a ratio of fluorescence of the area filled with PALs and fluorescence of soma (Medvedev et al., 2014; Plata et al., 2018). We found that the VF of PALs was significantly increased after CR. This finding suggests that the diet increases astrocyte presence in the synaptic microenvironment.

At the level of the astrocytic network, morphological plasticity can affect astrocyte coupling. Astrocytes make connections to their neighbors through homocellular gap-junctions permeable to ions and small molecules (Anders et al., 2014; Dermietzel et al., 1989). The role of gap-junctions in K^+^ spatial buffering has been suggested (Kofuji and Newman, 2004; Ma et al., 2016). Ca^2+^ events propagate in the astrocytic network through two major mechanisms – extracellular, involving neurotransmitter release, and intracellular, through the gap-junctions (Fujii et al., 2017; Semyanov, 2019). Gap-junction coupling has two possibilities for plasticity: (1) change in the number of connections and (2) change in the connection strength. Both types of plasticity can be estimated with monitoring fluorescent dye diffusion via gap-junctions (Anders et al., 2014; Plata et al., 2018). We found that astrocytic coupling decreased, but the strength of this coupling did not change significantly after CR. Notably, the level of connexin 43, which forms astrocytic gap-junctions was not affected in CR mice.

Astrocytes being secretory cells of the CNS (Verkhratsky et al., 2016) may release numerous factors affecting synaptogenesis (through thrombospondins) and synaptic remodeling (through supplying neurons for example with cholesterol) or supplying neurons with neurotransmitter precursors (glutamine and L-serine). Often the astrocytic secretion depends on Ca^2+^ activity in these cells (Araque et al., 2014; Zorec et al., 2012). In addition, Ca^2+^ activity itself regulates synaptic coverage by PALs (Tanaka et al., 2013). We found no significant difference in the frequency of Ca^2+^ events in CR mice. However, due to astrocyte uncoupling, the Ca^2+^ events could not propagate through the gap-junctions and became less spread. In contrast, the duration of Ca^2+^ events significantly increased, compensating for their smaller size. As a result, the integral of Ca^2+^ events (x-y-time) was not affected. Thus, overall Ca^2+^ activity was not increased or decreased but rather restructured in the spatiotemporal domain by CR.

The role of spatiotemporal patterns of astrocytic Ca^2+^ activity has recently started to be appreciated (Kanakov et al., 2019; Semyanov, 2008; Semyanov, 2019). Individual Ca^2+^ transients can trigger the local release of gliotransmitters (Zorec et al., 2012) or change the local synaptic microenvironment (e.g., by retracting or extending perisynaptic processes) (Tanaka et al., 2013). Thus, the spatiotemporal pattern of Ca^2+^ activity in the astrocytic network can form a ‘guiding template’ which shapes signal propagation and synaptic plasticity in the neuronal network. The decrease in the size of Ca^2+^ events observed after CR should increase the spatial resolution of the pattern. We can contemplate that more information can be encoded within a unit of volume in neuron-astrocytic network with such pattern. Prolongation of Ca^2+^ events may also be necessary to ensure that a sufficient number of synapses is affected within the active spot. Thus, shorter and longer Ca^2+^ events in the CR mice should influence more spatially compact groups of synapses. This may be beneficial for triggering of dendritic spikes that depend on co-activation of spatially colocalized synaptic inputs to the same dendric branch (Spruston, 2008; Varga et al., 2011). The dendritic spikes are required for LTP at synapses on hippocampal pyramidal neurons (Brandalise et al., 2016; Kim et al., 2015) and hippocampal granular cells (Kim et al., 2018). Thus, our findings provide an insight into how the information encoding and storage in the brain can become more efficient after CR.

PALs regulate glutamatergic transmission through active control over glutamate dynamics in the synaptic cleft (with the help of dedicated transporters), through K^+^ clearance and through supplying neurons with glutamine, an obligatory precursor of glutamate (by glutamate-GABA glutamine shuttle). Indeed, the increased presence of PALs in synaptic microenvironment correlated with the reduced spillover of both K^+^ and glutamate. The higher efficiency of glutamate uptake and, hence, lower ambient glutamate concentrations can be responsible for further functional changes in astrocytes. Extracellular elevations of glutamate positively regulate glutamine synthase (Lehmann et al., 2009). Lower extracellular concentrations of glutamate can possibly underlie a reduction in glutamine synthetase expression. Indeed, Western blot analysis revealed decreased expression of glutamine synthase in CR mice. This, in turn, can reduce glutamine synthesis which is required both for glutamatergic and GABAergic signaling in the brain (Rose et al., 2013).

We estimated the efficiency of glutamatergic signaling in hippocampal CA1 through astrocytic recordings of I_K_ and I_GluT_. The five-pulse ratio of I_K_ became higher after CR. Because most of I_K_ during synaptic transmission reflects its efflux through postsynaptic AMPA and NMDA receptors (Shih et al., 2013; Sibille et al., 2014), this finding suggests activity-dependent facilitation of synaptic transmission. However, the five-pulse ratio of I_GluT_ was not affected by CR, ruling out a modification of presynaptic release and highlighting postsynaptic mechanisms.

Next, we tested whether increased synaptic enwrapping by astrocytes and glutamate clearance correlates with enhanced synaptic plasticity (Valtcheva and Venance, 2019). The input-output characteristics of synaptic transmission (PrV vs. stimulus and fEPSP vs. stimulus) were not different in CR mice, suggesting that the diet did not influence baseline synaptic signaling. Consistent with previous reports efficient glutamate clearance by the astrocytes correlated with enhanced LTP in CR mice (Katagiri et al., 2001; Scimemi et al., 2009; Wang et al., 2006)

In conclusion, we find that caloric restriction evokes astroglial plasticity represented by an increased outgrowth of astrocytic perisynaptic processes and astrocytic presence in the synaptic microenvironment. This facilitates glutamate and K^+^ clearance and limits their spillover. In addition to morpho-functional remodeling of single astrocytes, the astrocytes in syncytia get uncoupled. This correlated with changes in the pattern of Ca^2+^ events in the astrocyte population: unable to propagate through the gap-junctions, Ca^2+^ events became smaller in size. However, the proportion of longer events increased, reflecting the outgrowth of astrocytic processes. All these events taken together affect neuronal networks and enhance neural plasticity required for adaptation to new feeding pattern.

## Methods

### Caloric restriction

The experiments were performed in two groups of mice from the age of 2 months. The animals of the first group were held in individual cages and received food *ad libitum* for a month (control). The amount of food was weighted, and 70% of the average daily consumptions per animal was fed to each animal in the second group (caloric restriction, CR).

### Slice preparation

All procedures were done in accordance with the University of Nizhny Novgorod regulations. The mice were anesthetized with isoflurane (1-chloro-2,2,2-trifluoroethyl-difluoromethyl ether) before being sacrificed. The brains were exposed and placed in ice-cold solution containing (in mM): 50 sucrose; 87 NaCl; 2.5 KCl; 8.48 MgSO4; 1.24 NaH2PO4; 26.2 NaHCO3; 0.5 CaCl2; 22 D-Glucose. Hippocampi dissected and cut in transverse slices (350 µm) using vibrating microtome (Microm HM650 V; Thermo Fisher Scientific). Slices were left to recover for 1 hour at 34° submerged in a solution containing (in mM): 119 NaCl, 2.5 KCl, 1.3 MgSO4, 1 NaH2PO4, 26.2 NaHCO3, 1 CaCl2, 1.6 MgCl2, 22 mM D-Glucose. The experiments were carried out at 34°C in immersion chambers with continuous perfusion (1-3 ml/min) by artificial cerebrospinal fluid (ACSF) containing (in mM): 119 NaCl; 2.5 KCl; 1.3 MgSO_4_; 1 NaH_2_PO_4_; 26.2 NaHCO_3_; 2 CaCl_2_; 11 D-Glucose. All solutions had an osmolarity of 300 ± 5 mOsm and a pH of 7.4 and were continuously bubbled with 95% O2 and 5% CO2. Hippocampal slices so prepared were used for both electrophysiological and imaging experiments.

### Field potential recording

Field excitatory postsynaptic potentials (fEPSPs) were recorded from CA1 *str.radiatum* using glass microelectrodes (2–5 MΩ) filled with normal Ringer solution. fEPSPs were evoked with extracellular stimulation of the Schaffer collaterals by using bipolar stimulating electrode (FHC, Bowdoinham, USA) placed in the *str.radiatum* at the CA1– CA2 border. The stimulation was performed with rectangular pulses (duration 0.1 ms every 20 s) with DS3 isolated current stimulator (Digitimer Ltd, UK). Responses were amplified using Multiclamp 700b amplifier (Molecular Devices, USA) and were digitized and recorded on a personal computer using ADC/DAC NI USB-6221 (National Instruments, USA), shielded connector block BNC2110 and WinWCP v5.2.3 software by John Dempster (University of Strathclyde). Electrophysiological data were analyzed with the Clampfit 10.2 program (Molecular Devices, USA).

The dependence of field response amplitude on stimulation strength was determined by increasing the current intensity from 100 to 400 μA. For each fEPSP, the amplitude and the slope of the rising phase at a level of 20–80% of the peak amplitude were measured. The presynaptic fiber volley (PrV) was quantified by the peak amplitude. The stimulus intensity was chosen from the amplitude of fEPSP in 40–50% of the amplitude where the population spike appeared for the first time. The strength of stimulation was unvaried during the experiments, usually being 100–150 μA.

The LTP induction was started only if the stable amplitude of the baseline fEPSP had been recorded for 15 min. Three trains of high-frequency stimulation (HFS, 100 pulses at 100 Hz, with an inter-train interval of 20 s protocol) was applied to induce LTP. The fEPSPs were recorded after induction protocol for 60 min. The baseline fEPSP and two types of the potentiated fEPSP (first recorded 0–10 min, second 40–50 min after HFS) averaged separately to measure LTP magnitude.

### Whole-cell electrophysiology recordings

Astrocytes were recorded in whole-cell voltage-clamp mode using Multiclamp 700B amplifier (Molecular Devices, USA). Borosilicate pipettes (3 - 5 MΩ) were filled with an internal solution containing (in mM): 135 KCH_3_SO_3_, 10 HEPES, 10 Na_2_-phosphocreatine, 4 MgCl_2_, 4 Na_2_-ATP, 0.4 Na-GTP (pH adjusted to 7.2 with KOH; osmolarity to 295 mOsm). 50 μM Alexa Fluor 594 (Invitrogen, USA) was added to the internal solution for morphology study.

Bipolar extracellular tungsten electrodes (FHC, Bowdoinham, USA) were placed in *str.radiatum* between CA1 and CA3 areas. Astrocytes were chosen in the *str.radiatum* at 100 – 200 µm away from the stimulating electrode. The cells were voltage-clamped at −80 mV. Passive astrocytes were identified by their small soma (5 – 10 µm diameter), low resting membrane potential (~ 80 mV), low input resistance (< 20 MΩ) and linear current-voltage relationship (IV-curve). Cycles of one, four and five electrical stimuli (50 Hz) were applied to Schaffer collaterals to induce synaptic currents in the astrocytes followed by a voltage step of −5 mV for monitoring cell input resistance. Signals were sampled at 5 kHz and filtered at 2 kHz and subsequently analyzed using custom-written MATLAB scrips (MathWorks, USA). Astrocyte currents in response to single, four and five stimuli were baseline subtracted and averaged correspondingly. Current in response to the fifth stimulus was obtained by the subtraction of current evoked by four stimuli from current evoked by five stimuli.

The membrane currents produced by synaptic stimulation in astrocytes represent a superposition of several components of which the main are glutamate transporter current (I_GluT_) and K^+^ inward current (I_K_) (Shih et al., 2013; Sibille et al., 2014).

### Analysis of potassium current (I_K_)

I_K_ is the slowest current in astrocytes induced by synaptic simulation and lasts for hundreds of ms. Without using any pharmacological intervention, I_K_ was analyzed 200 ms after the last stimulus. At this time point, I_K_ is not contaminated by the current mediated by field potential and I_GluT_. If the peak of I_K_ was not found after 200 ms, the I_K_ value at 200 ms considered as I_K_ amplitude. The decay was fitted with mono-exponential function and τ_decay_ calculated.

### Analysis of glutamate transporter current (I_GlutT_)

I_GluT_ was obtained in the presence of 25 µM NBQX, 50 µM D-AP5 and 100 µM picrotoxin (all from Tocris Bioscience, UK) which blocked AMPA, NMDA and GABA_A_ receptors, respectively. After obtaining 10 to 20 recordings for each cycle, an excitatory amino acid transporter (EAATs) blocker DL-TBOA (100 µM, Tocris Bioscience, UK) was added to the bath. Pure I_K_ was then recorded in response to a single stimulus. This I_K_ was mathematically reconstructed with two mono-exponential fits of its rise and decay segments. The resultant mathematical noiseless wave was subtracted from synaptically induced combined currents to obtain pure I_GluT_.

### Single-cell morphometry

Astrocytes were loaded with fluorescent dye Alexa Fluor 594 through the patch pipette which was used for electrophysiology recordings. The cells were visualized with Axio Examiner Z1 (Zeiss) microscope equipped with infrared differential interference contrast (IR-DIC). Images were collected with W Plan-APOCHROMAT 40×/1.0 water immersion objective, using the Zeiss LSM 7 MP system (Carl Zeiss, Germany) coupled with Ti:sapphire broadband laser multi-photon system Chameleon Vision II (Coherent, UK).

### Sholl analysis

All processing steps were performed using image-funcut library [image-funcut, https://github.com/abrazhe/image-funcut] and other custom-written Python scripts, using Scikit-Image [scikit, http://scikit-image.org/] and Sci-Py [scipy, http://www.scipy.org/] libraries. In brief, z-stacks corresponding to the emission spectrum (565-610 nm) of Alexa Fluor 594 were re-sampled to the same lateral resolution of 0.25 µm/px. Coherence-enhancing diffusion filtering was performed for each image in the stack to enhance all filamentous structures. Then the z-stack was collapsed with maximal intensity projection along the z-axis, and adaptive threshold filtering was applied to create the binary mask. Sholl profile was acquired automatically as a number of intersections of circles with increasing radii from the center of the astrocyte soma.

### Volume fraction (VF) estimation

The VF of astrocyte fine processes was estimated as previously described (Medvedev et al., 2014; Plata et al., 2018; Wu et al., 2019). The image containing the middle of astrocyte soma was selected in the z-stack. Special attention was paid that the fluorescence of soma was not saturated. Eight radial lines were plotted from the center of soma at the angle of 22.5° from each other. The fluorescent profiles along these lines were plotted, and large fluctuations (> 10%) of fluorescence corresponding to astrocytic branches were cut out. The mean fluorescence profile was calculated for each cell. The fluorescence profile values were divided by the peak fluorescence in soma to obtain the VF estimation:

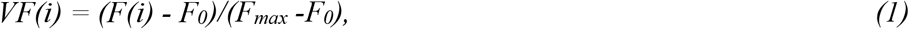

where *VF(i)* is VF of a particular point in the profile*, F(i)* - the fluorescence of this point, *F*_*max*_ - the highest fluorescence value of the line profile in the soma, *F*_*0*_ - the background fluorescence. *F*_*0*_ is the mean fluorescence intensity of a circle (diameter 10 µm) outside the astrocytic arbor without any stained structures. The VF is presented for a segment between 8 and 30 µm to exclude the soma and bias due to the asymmetry of the astrocyte domain. The values of mean VF presented in the text calculated from the mean VF for each astrocyte profile.

### Astrocyte coupling analysis

The astrocyte gap-junction coupling was estimated through fluorescent dye diffusion as previously described (Anders et al., 2014; Plata et al., 2018). A single z-stack of images with dimensions of × = 200 µm, y = 200 μm, z = 70 µm was obtained 25 min after whole-cell establishment. Averaged somatic fluorescence intensities of the coupled cells were normalized to the soma fluorescence of the patched astrocyte. The distance to the patched astrocyte was calculated in 3D using the Pythagorean theorem. The relationship between the distance and normalized fluorescence was fitted with a monoexponential function to obtain coupling length constant (C_λ_):

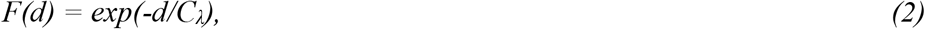

where *F(d)* is normalized fluorescence of coupled astrocyte soma at the distance *d* from the patched astrocyte.

### Ca^2+^ imaging

Astrocyte Ca^2+^ activity was recorded with a confocal microscope, Zeiss LSM DuoScan 510, in CA1 *str.radiatum* of acute hippocampal slices pre-incubated with Ca^2+^ dye, Oregon Green 488 BAPTA-1 AM (OGB, Invitrogen, USA) and an astrocyte-specific marker, sulforhodamine 101 (100 nM, Invitrogen, USA). After the preparation, the slices were transferred to a 3 ml incubation chamber with constantly bubbled ACSF containing both dyes. OGB was initially dissolved to 0.795 mM in 0.8% Pluronic F-127 in DMSO. Then 3 µl of the dye was added to the chamber. After incubation for 40 - 45 min at 37 °C in the dark, the slices were transferred to the recording/imaging chamber for time-lapse imaging (one frame/s). OGB was excited with a 488 nm argon laser and imaged with an emission band-pass filter 500 - 530 nm; sulforhodamine 101 was excited with a 543 nm HeNe laser and imaged with an emission band-pass filter 650 - 710 nm. The imaging was performed for 10 min at 34 °C in normal ASCF, then 30 dark noise images were recorded.

The raw imaging data were exported to MATLAB. The median of the dark noise was calculated for each pixel and subtracted from the corresponding pixel intensity value of the fluorescence images. Then denoising was done with the BM3D algorithm (Danielyan et al., 2014). The movement artifacts were corrected with the single-step DFT algorithm (Guizar-Sicairos et al., 2008). The Ca^2+^ events (x-y-time, 3D Ca^2+^ signals) were detected with the adapted algorithm which we described previously (Plata et al., 2018; Wu et al., 2014). For each Ca^2+^ event the maximal projection (size), the duration and the integral were calculated.

### Western blotting

The hippocampi of CR and control mice were frozen in liquid nitrogen. Then each hippocampus was homogenized in RIPA buffer with SIGMAFAST protease inhibitor cocktail (Sigma-Aldrich, St. Louis, USA), diluted in loading buffer, submitted to gel electrophoresis, and blotted onto nitrocellulose membranes (GE Healthcare, Chicago, USA). The membranes were blocked in 5% BSA overnight and incubated for 1 hour with primary mouse antibodies against glutamine synthetase (GS, Sigma-Aldrich, MAB302), connexin 43 (Cx43, Santa-Cruz, sc-271837, Dallas, USA) or β-actin (R&D Systems, MAB8929, Minneapolis, USA). For detection of glutamate transporter 1 (GLT-1), primary rabbit antibodies were used (Abcam, 106289, Waltham, USA). Then membranes were rinsed in Tris-Buffer (pH =7.4) with 0.1 % of Tween-20 and incubated with HRP-conjugated secondary antibodies – anti-mouse IgG (Jackson Immunoresearch, 715-005-150, Ely, UK) or with anti-rabbit IGG (Abcam, 6721) for 1 hour. ECL substrate (Bio-Rad, Hercules, USA) was used for signal detection. Protein bands were visualized using a VersaDoc 4000 chemidocumenter (Bio-Rad). The intensity of protein bands was quantified using gel analyzer option of ImageJ software (NIH, Bethesda, USA). Statistical analysis was performed by GraphPad Prism v 6.0 (GraphPad Software, San Diego, USA).

### Statistical analysis

All data are presented as the mean ± standard error of the mean (SEM) except for the measurements of astrocytic Ca^2+^ events which followed a skewed distribution. Statistical significance was assessed using parametric Student’s *t*-test, nonparametric Mann-Whitney test and repeated measures two-way ANOVA as stated in the text. *p* < 0.05 was considered statistically significant.

## Acknowledgments

The work was supported by Volkswagen Stiftung research grant A115105 to AS and AV and COMFI grant 17-00-00412 (K) from RFBR for joint research of AS (grant 17-00-00409) and AB (grant 17-00-00407)

## References

Anders, S., Minge, D., Griemsmann, S., Herde, M.K., Steinhauser, C., and Henneberger, C. (2014). Spatial properties of astrocyte gap junction coupling in the rat hippocampus. Philos Trans R Soc Lond B Biol Sci 369, 20130600.

Araque, A., Carmignoto, G., Haydon, P.G., Oliet, S.H., Robitaille, R., and Volterra, A. (2014). Gliotransmitters Travel in Time and Space. Neuron 81, 728–739.

Brandalise, F., Carta, S., Helmchen, F., Lisman, J., and Gerber, U. (2016). Dendritic NMDA spikes are necessary for timing-dependent associative LTP in CA3 pyramidal cells. Nat Commun 7, 13480–13480.

Brasnjo, G., and Otis, T.S. (2001). Neuronal Glutamate Transporters Control Activation of Postsynaptic Metabotropic Glutamate Receptors and Influence Cerebellar Long-Term Depression. Neuron 31, 607–616.

Castiglioni Jr, A.J., Legare, M.E., Busbee, D.L., and Tiffany-Castiglioni, E. (1991). Morphological changes in astrocytes of aging mice fed normal or caloric restricted diets. Age 14, 102–106.

Danielyan, A., Wu, Y.-W., Shih, P.-Y., Dembitskaya, Y., and Semyanov, A. (2014). Denoising of two-photon fluorescence images with Block-Matching 3D filtering. Methods 68, 308–316.

Dermietzel, R., Traub, O., Hwang, T.K., Beyer, E., Bennett, M.V., Spray, D.C., and Willecke, K. (1989). Differential expression of three gap junction proteins in developing and mature brain tissues. Proceedings of the National Academy of Sciences 86, 10148–10152.

Fernandez, B., Suarez, I., and Gonzalez, G. (1984). Topographical distribution of the astrocytic lamellae in the hypothalamus. Anat Anz 156, 31–37.

Fontan-Lozano, A., Lopez-Lluch, G., Delgado-Garcia, J.M., Navas, P., and Carrion, A.M. (2008). Molecular bases of caloric restriction regulation of neuronal synaptic plasticity. Mol Neurobiol 38, 167–177.

Fontana, L., Partridge, L., and Longo, V.D. (2010). Extending healthy life span--from yeast to humans. Science 328, 321–326.

Fujii, Y., Maekawa, S., and Morita, M. (2017). Astrocyte calcium waves propagate proximally by gap junction and distally by extracellular diffusion of ATP released from volume-regulated anion channels. Sci Rep 7, 13115.

Garcia-Caceres, C., Balland, E., Prevot, V., Luquet, S., Woods, S.C., Koch, M., Horvath, T.L., Yi, C.X., Chowen, J.A., Verkhratsky, A., et al. (2019). Role of astrocytes, microglia, and tanycytes in brain control of systemic metabolism. Nat Neurosci 22, 7–14.

Gavrilov, N., Golyagina, I., Brazhe, A., Scimemi, A., Turlapov, V., and Semyanov, A. (2018). Astrocytic Coverage of Dendritic Spines, Dendritic Shafts, and Axonal Boutons in Hippocampal Neuropil. Frontiers in cellular neuroscience 12.

Ge, W.P., and Duan, S. (2007). Persistent enhancement of neuron-glia signaling mediated by increased extracellular K+ accompanying long-term synaptic potentiation. J Neurophysiol 97, 2564–2569.

Guizar-Sicairos, M., Thurman, S.T., and Fienup, J.R. (2008). Efficient subpixel image registration algorithms. Opt Lett 33, 156–158.

Henneberger, C., Papouin, T., Oliet, S.H., and Rusakov, D.A. (2010). Long-term potentiation depends on release of D-serine from astrocytes. Nature 463, 232–236.

Kanakov, O., Gordleeva, S., Ermolaeva, A., Jalan, S., and Zaikin, A. (2019). Astrocyte-induced positive integrated information in neuron-astrocyte ensembles. Physical Review E 99, 012418.

Katagiri, H., Tanaka, K., and Manabe, T. (2001). Requirement of appropriate glutamate concentrations in the synaptic cleft for hippocampal LTP induction. European Journal of Neuroscience 14, 547–553.

Khakh, B.S., and Sofroniew, M.V. (2015). Diversity of astrocyte functions and phenotypes in neural circuits. Nat Neurosci 18, 942–952.

Kim, S., Kim, Y., Lee, S.-H., and Ho, W.-K. (2018). Dendritic spikes in hippocampal granule cells are necessary for long-term potentiation at the perforant path synapse. eLife 7, e35269.

Kim, Y., Hsu, C.-L., Cembrowski, M.S., Mensh, B.D., and Spruston, N. (2015). Dendritic sodium spikes are required for long-term potentiation at distal synapses on hippocampal pyramidal neurons. eLife 4, e06414.

Kofuji, P., and Newman, E.A. (2004). Potassium buffering in the central nervous system. Neuroscience 129, 1045–1056.

Kuhla, A., Lange, S., Holzmann, C., Maass, F., Petersen, J., Vollmar, B., and Wree, A. (2013). Lifelong caloric restriction increases working memory in mice. PLoS One 8, e68778.

Kullmann, D.M., and Asztely, F. (1998). Extrasynaptic glutamate spillover in the hippocampus: evidence and implications. Trends Neurosci 21, 8–14.

Lebedeva, A., Plata, A., Nosova, O., Tyurikova, O., and Semyanov, A. (2018). Activity-dependent changes in transporter and potassium currents in hippocampal astrocytes. Brain research bulletin 136, 37–43.

Lehmann, C., Bette, S., and Engele, J. (2009). High extracellular glutamate modulates expression of glutamate transporters and glutamine synthetase in cultured astrocytes. Brain Research 1297, 1–8.

Lin, A.L., Coman, D., Jiang, L., Rothman, D.L., and Hyder, F. (2014). Caloric restriction impedes age-related decline of mitochondrial function and neuronal activity. J Cereb Blood Flow Metab 34, 1440–1443.

Ma, B., Buckalew, R., Du, Y., Kiyoshi, C.M., Alford, C.C., Wang, W., McTigue, D.M., Enyeart, J.J., Terman, D., and Zhou, M. (2016). Gap junction coupling confers isopotentiality on astrocyte syncytium. Glia 64, 214–226.

Maalouf, M., Rho, J.M., and Mattson, M.P. (2009). The neuroprotective properties of calorie restriction, the ketogenic diet, and ketone bodies. Brain Res Rev 59, 293–315.

Magistretti, P.J. (2009). Neuroscience. Low-cost travel in neurons. Science 325, 1349–1351.

Masoro, E.J. (2009). Caloric restriction-induced life extension of rats and mice: a critique of proposed mechanisms. Biochim Biophys Acta 1790, 1040–1048.

Massey, P.V., Johnson, B.E., Moult, P.R., Auberson, Y.P., Brown, M.W., Molnar, E., Collingridge, G.L., and Bashir, Z.I. (2004). Differential Roles of NR2A and NR2B-Containing NMDA Receptors in Cortical Long-Term Potentiation and Long-Term Depression. The Journal of Neuroscience 24, 7821–7828.

Mattison, J.A., Colman, R.J., Beasley, T.M., Allison, D.B., Kemnitz, J.W., Roth, G.S., Ingram, D.K., Weindruch, R., de Cabo, R., and Anderson, R.M. (2017). Caloric restriction improves health and survival of rhesus monkeys. Nat Commun 8, 14063.

Mattson, M.P. (2012). Energy intake and exercise as determinants of brain health and vulnerability to injury and disease. Cell Metab 16, 706–722.

Mattson, M.P. (2015). Lifelong brain health is a lifelong challenge: from evolutionary principles to empirical evidence. Ageing Res Rev 20, 37–45.

McCay, C.M., Crowell, M.F., and Maynard, L.A. (1935). The effect of retarded growth upon the lenght of life span and upon the ultimate body size. J Nutr 10, 63–79.

Medvedev, N., Popov, V., Henneberger, C., Kraev, I., Rusakov, D.A., and Stewart, M.G. (2014). Glia selectively approach synapses on thin dendritic spines. Philos Trans R Soc Lond B Biol Sci 369, 20140047.

Monai, H., Ohkura, M., Tanaka, M., Oe, Y., Konno, A., Hirai, H., Mikoshiba, K., Itohara, S., Nakai, J., Iwai, Y., and Hirase, H. (2016). Calcium imaging reveals glial involvement in transcranial direct current stimulation-induced plasticity in mouse brain. Nat Commun 7, 11100.

Murphy, T., Dias, G.P., and Thuret, S. (2014). Effects of diet on brain plasticity in animal and human studies: mind the gap. Neural Plast 2014, 563160.

Nedergaard, M., and Verkhratsky, A. (2012). Artifact versus reality - how astrocytes contribute to synaptic events. Glia 60, 1013–1023.

Ngandu, T., Lehtisalo, J., Solomon, A., Levalahti, E., Ahtiluoto, S., Antikainen, R., Backman, L., Hanninen, T., Jula, A., Laatikainen, T., et al. (2015). A 2 year multidomain intervention of diet, exercise, cognitive training, and vascular risk monitoring versus control to prevent cognitive decline in at-risk elderly people (FINGER): a randomised controlled trial. Lancet 385, 2255–2263.

Okada, M., Nakanishi, H., Amamoto, T., Urae, R., Ando, S., Yazawa, K., and Fujiwara, M. (2003). How does prolonged caloric restriction ameliorate age-related impairment of long-term potentiation in the hippocampus? Brain Res Mol Brain Res 111, 175–181.

Oliet, S.H., Piet, R., and Poulain, D.A. (2001). Control of glutamate clearance and synaptic efficacy by glial coverage of neurons. Science 292, 923–926.

Patrushev, I., Gavrilov, N., Turlapov, V., and Semyanov, A. (2013). Subcellular location of astrocytic calcium stores favors extrasynaptic neuron-astrocyte communication. Cell calcium 54, 343–349.

Pellerin, L., and Magistretti, P.J. (2012). Sweet sixteen for ANLS. J Cereb Blood Flow Metab 32, 1152–1166.

Perez-Alvarez, A., Navarrete, M., Covelo, A., Martin, E.D., and Araque, A. (2014). Structural and Functional Plasticity of Astrocyte Processes and Dendritic Spine Interactions. The Journal of Neuroscience 34, 12738–12744.

Pifferi, F., Terrien, J., Marchal, J., Dal-Pan, A., Djelti, F., Hardy, I., Chahory, S., Cordonnier, N., Desquilbet, L., Hurion, M., et al. (2018). Caloric restriction increases lifespan but affects brain integrity in grey mouse lemur primates. Commun Biol 1, 30.

Plata, A., Lebedeva, A., Denisov, P., Nosova, O., Postnikova, T.Y., Pimashkin, A., Brazhe, A., Zaitsev, A.V., Rusakov, D.A., and Semyanov, A. (2018). Astrocytic Atrophy Following Status Epilepticus Parallels Reduced Ca2+ Activity and Impaired Synaptic Plasticity in the Rat Hippocampus. Frontiers in Molecular Neuroscience 11.

Redman, L.M., Martin, C.K., Williamson, D.A., and Ravussin, E. (2008). Effect of caloric restriction in non-obese humans on physiological, psychological and behavioral outcomes. Physiol Behav 94, 643–648.

Ribeiro, L.C., Quincozes-Santos, A., Leite, M.C., Abib, R.T., Kleinkauf-Rocha, J., Biasibetti, R., Rotta, L.N., Wofchuk, S.T., Perry, M.L.S., Gonçalves, C.-A., and Gottfried, C. (2009). Caloric restriction increases hippocampal glutamate uptake and glutamine synthetase activity in Wistar rats. Neuroscience Research 64, 330–334.

Romanos, J., Benke, D., Saab, A.S., Zeilhofer, H.U., and Santello, M. (2019). Differences in glutamate uptake between cortical regions impact neuronal NMDA receptor activation. Communications Biology 2, 127.

Rose, Christopher F., Verkhratsky, A., and Parpura, V. (2013). Astrocyte glutamine synthetase: pivotal in health and disease. Biochemical Society Transactions 41, 1518–1524.

Rusakov, D.A., and Kullmann, D.M. (1998). Extrasynaptic glutamate diffusion in the hippocampus: ultrastructural constraints, uptake, and receptor activation. J Neurosci 18, 3158–3170.

Schiweck, J., Eickholt, B.J., and Murk, K. (2018). Important Shapeshifter: Mechanisms Allowing Astrocytes to Respond to the Changing Nervous System During Development, Injury and Disease. Frontiers in cellular neuroscience 12, 261–261.

Scimemi, A., and Diamond, J.S. (2013). Deriving the time course of glutamate clearance with a deconvolution analysis of astrocytic transporter currents. J Vis Exp.

Scimemi, A., Tian, H., and Diamond, J.S. (2009). Neuronal transporters regulate glutamate clearance, NMDA receptor activation, and synaptic plasticity in the hippocampus. The Journal of neuroscience: the official journal of the Society for Neuroscience 29, 14581–14595.

Semyanov, A. (2008). Can diffuse extrasynaptic signaling form a guiding template? Neurochemistry International 52, 31–33.

Semyanov, A. (2019). Spatiotemporal pattern of calcium activity in astrocytic network. Cell Calcium 78, 15–25.

Shih, P.-Y., Savtchenko, L.P., Kamasawa, N., Dembitskaya, Y., McHugh, T.J., Rusakov, D.A., Shigemoto, R., and Semyanov, A. (2013). Retrograde Synaptic Signaling Mediated by K+ Efflux through Postsynaptic NMDA Receptors. Cell reports 5, 941–951.

Sibille, J., Pannasch, U., and Rouach, N. (2014). Astroglial potassium clearance contributes to short-term plasticity of synaptically evoked currents at the tripartite synapse. J Physiol 592, 87–102.

Song, I.Y., Song, M., T., T., and Lee, H. (2018). The landscape of smart aging: Topics, applications, and agenda. Data & Knowledge Engineering 115, 68–79.

Spolidoro, M., Baroncelli, L., Putignano, E., Maya-Vetencourt, J.F., Viegi, A., and Maffei, L. (2011). Food restriction enhances visual cortex plasticity in adulthood. Nat Commun 2, 320.

Spruston, N. (2008). Pyramidal neurons: dendritic structure and synaptic integration. Nat Rev Neurosci 9, 206–221.

Tanaka, M., Shih, P.-Y., Gomi, H., Yoshida, T., Nakai, J., Ando, R., Furuichi, T., Mikoshiba, K., Semyanov, A., and Itohara, S. (2013). Astrocytic Ca2+ signals are required for the functional integrity of tripartite synapses. Molecular brain 6.

Valtcheva, S., and Venance, L. (2019). Control of Long-Term Plasticity by Glutamate Transporters. Frontiers in Synaptic Neuroscience 11.

Varga, Z., Jia, H., Sakmann, B., and Konnerth, A. (2011). Dendritic coding of multiple sensory inputs in single cortical neurons in vivo. Proceedings of the National Academy of Sciences 108, 15420–15425.

Veech, R.L., Bradshaw, P.C., Clarke, K., Curtis, W., Pawlosky, R., and King, M.T. (2017). Ketone bodies mimic the life span extending properties of caloric restriction. IUBMB Life 69, 305–314.

Verkhratsky, A., Matteoli, M., Parpura, V., Mothet, J.P., and Zorec, R. (2016). Astrocytes as secretory cells of the central nervous system: idiosyncrasies of vesicular secretion. EMBO J 35, 239–257.

Verkhratsky, A., and Nedergaard, M. (2014). Astroglial cradle in the life of the synapse. Philos Trans R Soc Lond B Biol Sci 369, 20130595.

Verkhratsky, A., and Nedergaard, M. (2018). Physiology of Astroglia. Physiol Rev 98, 239–389.

Wang, Z.-Y., Zhang, Y.-Q., and Zhao, Z.-Q. (2006). Inhibition of tetanically sciatic stimulation-induced LTP of spinal neurons and Fos expression by disrupting glutamate transporter GLT-1. Neuropharmacology 51, 764–772.

Wong, T.P., Howland, J.G., Robillard, J.M., Ge, Y., Yu, W., Titterness, A.K., Brebner, K., Liu, L., Weinberg, J., Christie, B.R., et al. (2007). Hippocampal long-term depression mediates acute stress-induced spatial memory retrieval impairment. Proceedings of the National Academy of Sciences 104, 11471–11476.

Wu, Y.-W., Gordleeva, S., Tang, X., Shih, P.-Y., Dembitskaya, Y., and Semyanov, A. (2019). Morphological profile determines the frequency of spontaneous calcium events in astrocytic processes. Glia 67, 246–262.

Wu, Y.-W., Tang, X., Arizono, M., Bannai, H., Shih, P.-Y., Dembitskaya, Y., Kazantsev, V., Tanaka, M., Itohara, S., Mikoshiba, K., and Semyanov, A. (2014). Spatiotemporal calcium dynamics in single astrocytes and its modulation by neuronal activity. Cell calcium 55, 119–129.

Yang, J., Wang, Q., He, F., Ding, Y., Sun, Q., Hua, T., and Xi, M. (2016). Dietary Restriction Affects Neuronal Response Property and GABA Synthesis in the Primary Visual Cortex. PLoS One 11, e0149004.

Zeltser, L.M., Seeley, R.J., and Tschop, M.H. (2012). Synaptic plasticity in neuronal circuits regulating energy balance. Nat Neurosci 15, 1336–1342.

Zorec, R., Araque, A., Carmignoto, G., Haydon, P.G., Verkhratsky, A., and Parpura, V. (2012). Astroglial excitability and gliotransmission: an appraisal of Ca2+ as a signalling route. ASN neuro 4.

