## Supporting information for "An astrocytic basis of caloric restriction action on the brain plasticity"

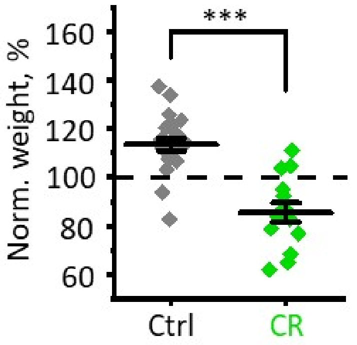


**Figure S1. CR reduces mouse weight**

2-month-old mice were separated into two groups. All animals were kept in solitary cages. The mice of the first group received food *ad libitum* - i.e., control group (Ctrl). The mice in the second group received 70% of the mean food consumption of the mice in the first group – i.e., CR group. After one month of the diet, the control mice gained weight to 114 ± 3 % ($n$ = 22, $p < 0.001$, one sample $t$-test), while the animals in CR group lost weight to 85 ± 4 % ($n$ = 14; $p < 0.001$, one sample $t$-test; $p < 0.001$, two sample $t$-test for comparison between two groups)

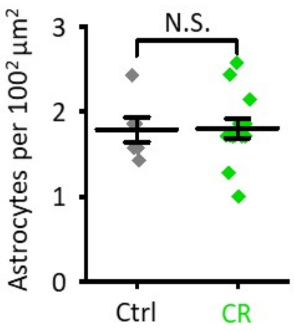


**Figure S2. The absence of significant changes in astrocyte density in CR**

Astrocytes labeled with specific marker sulforhodamine 101 were counted. No significant difference in the astrocyte density was observed in the hippocampus of control (grey diamonds) and of CR (green diamonds) mice. The data are presented as mean ± SEM. N.S. $p > 0.05$, two sample *t*-test.

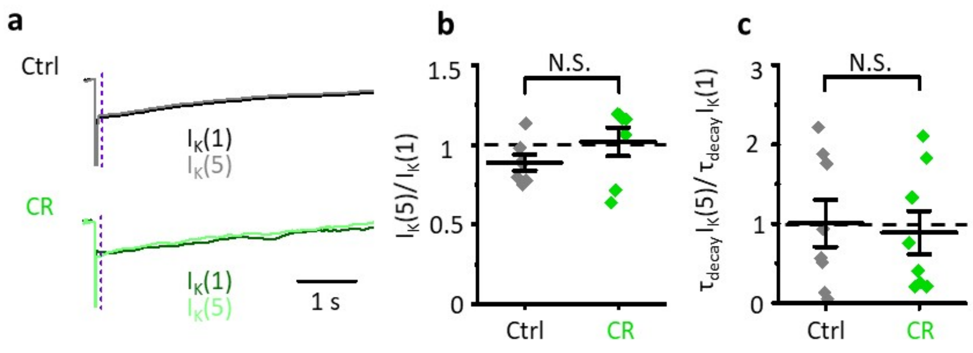


**Figure S3. $I_K$ in the presence of NMDA, AMPA and $GABA_A$ receptor blockers**

**a.** Representative current traces to a single stimulus (dark traces) and the fifth stimulus (light traces) in control (Ctrl, grey) and CR mice (green). The dashed line shows where the measurement of $I_K(1)$ and $I_K(5)$ amplitudes and decay times were taken to avoid the interference of $I_{GluT}$. **b.** The summary plot showing no significant difference in the five pulse ratio – i.e., $I_K(5)/I_K(1)$, between control (grey diamonds) and CR (green diamonds) mice. **c.** The summary plot showing no difference in the five pulse ratio of decay time – i.e., $\tau_{decay}$ $I_K(5)/\tau_{decay}$ $I_K(1)$, between control (grey diamonds) and CR (green diamonds) mice.

The data are presented as mean ± SEM; N.S. $p > 0.05$, two sample *t*-test.
